# Regulation of host metabolism and defense strategies to survive neonatal infection

**DOI:** 10.1101/2024.02.23.581534

**Authors:** Ziyuan Wu, Nguyen Tran Nam Tien, Ole Bæk, Jingren Zhong, Björn Klabunde, Tinh Thu Nguyen, Nguyen Thi Hai Yen, Nguyen Phuoc Long, Duc Ninh Nguyen

## Abstract

Two distinct defense strategies, resistance and tolerance, enable a host to survive infectious diseases. Newborns, constrained by limited energy reserves, predominantly rely on tolerance to cope with infection. However, this approach may fail as pathogen levels surpass a critical threshold, prompting a shift to resistance that can lead to dysregulated immune responses and sepsis. The mechanisms governing the interplay between tolerance and resistance in newborns remain poorly understood. Here, we compare metabolic traits and defense strategies between survivors and non-survivors in *Staphylococcus epidermidis* (*S. epidermidis*)-infected preterm piglets, mimicking infection in preterm infants. Relative to non-survivors, survivors displayed elevated resistance during the early phase of infection, followed by stronger tolerance in later stages. Conversely, animals succumbing to sepsis showed clear signs of respiratory and metabolic acidosis, together with exaggerated inflammation and organ dysfunctions. Hepatic transcriptomics revealed a strong association between the tolerance phenotype and heightened oxidative phosphorylation in survivors, coupled with suppressed glycolysis and immune signaling. Plasma metabolomics supported the finding of enhanced mitochondrial metabolism in survivors. Our findings suggest a link between mitochondrial metabolism, disease tolerance, and ultimately improved survival during infections in newborns. Metabolic regulations related to tolerance may be exploited to discover novel therapeutics for neonatal infection.

**Conflict-of-interest statement:** The authors have declared that no conflict of interest exists.

## Introduction

Early life infection resulting in neonatal sepsis is one of the leading causes of death in newborns, especially those born prematurely (1–3). The majority of clinical complications during neonatal sepsis are not derived directly from the pathogens but from the dysregulated immune responses and subsequent tissue damage (4). Shortly after preterm birth, newborn hosts have lower levels of maternally transferred immunoglobulin G (IgG), and circulating immune cells with reduced capacities to produce inflammatory cytokines (5). However, even with these compromises, their immune system can still trigger dysregulated immune responses upon infection with excessive inflammation, leading to sepsis, tissue damage, and a high risk of death (6).

Two distinct defense strategies against pathogens have been described in mammals (7, 8): resistance, which entails active immune responses to eradicate the invading microbe; and tolerance, which promotes host health while not eliminating pathogens to minimize immunopathological damage. Resistance and tolerance are two separate but complementary host defense strategies that can be switched during infection. Newborns heavily rely on tolerance strategy, thus decreasing the susceptibility to tissue damage from active resistance (4, 9). Specifically, newborns demonstrate an impressive ability to tolerate circulating bacterial loads that are 10-100 times higher than those tolerated by adults (3, 9, 10). However, once a certain threshold of bacterial loads is reached, a resistance response becomes predominant, often with dysregulated inflammation and tissue damage, explaining why newborns are more prone to sepsis. This delicate balance highlights a reliance on tolerance and a higher threshold for active resistance in neonates. Neonatal tolerance plays a crucial role in ensuring survival following extrauterine exposure after birth and might be mediated by the unique metabolic state during the immediate postnatal period. Specifically, neonates are born with low levels of fat, glycogen, and muscle mass, while the energetic demands for growth and development necessitate maximal energy expenditure even at rest (5, 11). Energy-intensive resistance-type infection responses may be minimized in favor of tolerance to conserve resources, reflecting a strategic adaptation to their limited energy reserves (4).

Sepsis is currently recognized to involve extensive host metabolic changes. The paradigm shift of defense strategies underscores the intricate interplay between metabolism and immunity. Upon Toll-like receptor (TLR) stimulation, immune effector cells (granulocytes, M1 macrophages, cytotoxic and helper T-cells, and NK cells) activate resistance responses and shift to aerobic glycolysis, necessitating a continuous glucose supply to maintain inflammatory responses (12, 13). This heightened energy demand for resistance compels infected neonates to divert resources from essential growth and development, thereby increasing susceptibility to organ failure due to energy deficits (4). Conversely, immune regulatory cells (Tregs, M2 macrophages, and memory T cells) involved in the tolerance response, adapt by increasing the uptake of exogenous fatty acids and sustaining elevated levels of β-oxidation and oxidative phosphorylation (OXPHOS) (12, 13). This adaptation is crucial during infection and sepsis, particularly in newborns, whose unique metabolic demands make these metabolic components of the immune response more critical to their survival. We have previously developed a *S. epidermidis*-infected preterm piglets model, mimicking infected preterm infants (14–17). We have demonstrated that high parenteral glucose supply exaggerates hepatic aerobic glycolysis and systemic inflammation that ultimately leads to sepsis (14, 17). This underscores the critical role of energy metabolism in the progression and outcome of sepsis. Here, we investigate how the regulation of resistance and tolerance mechanisms interplay with host energy metabolism. We identify that infected survivors of sepsis show enhanced resistance responses at the early stage of infection and a stronger tolerance strategy later than those non-survivors. This is associated with a superior capability to modulate infection-induced aerobic glycolysis and inflammation alongside augmented energy production at the endpoint. Our study provides a significant advancement in understanding the interplay between sepsis pathogenesis and host adaptive responses, particularly tolerance, and may guide future strategies to improve tolerance as a potential target against neonatal sepsis.

## Results

### Clinical response disparities between infected survivors and non-survivors

Using *S.epidermidis* infected preterm piglets as a model for septic preterm infants, we explored the clinical, cellular, and molecular responses to *S. epidermidis* infection over 15 h post-inoculation (Fig. 1A). After 15 h, 56% of piglets (15/27 infected piglets) developed sepsis, predefined as arterial blood pH of ≤ 7.1 and clinical signs of deep lethargy, respiratory arrest, and skin bleeding (17). This allowed for the division of piglets into three groups: survival (n = 12), non-survival (n = 15), and uninfected (n = 9).

**Fig. 1:**
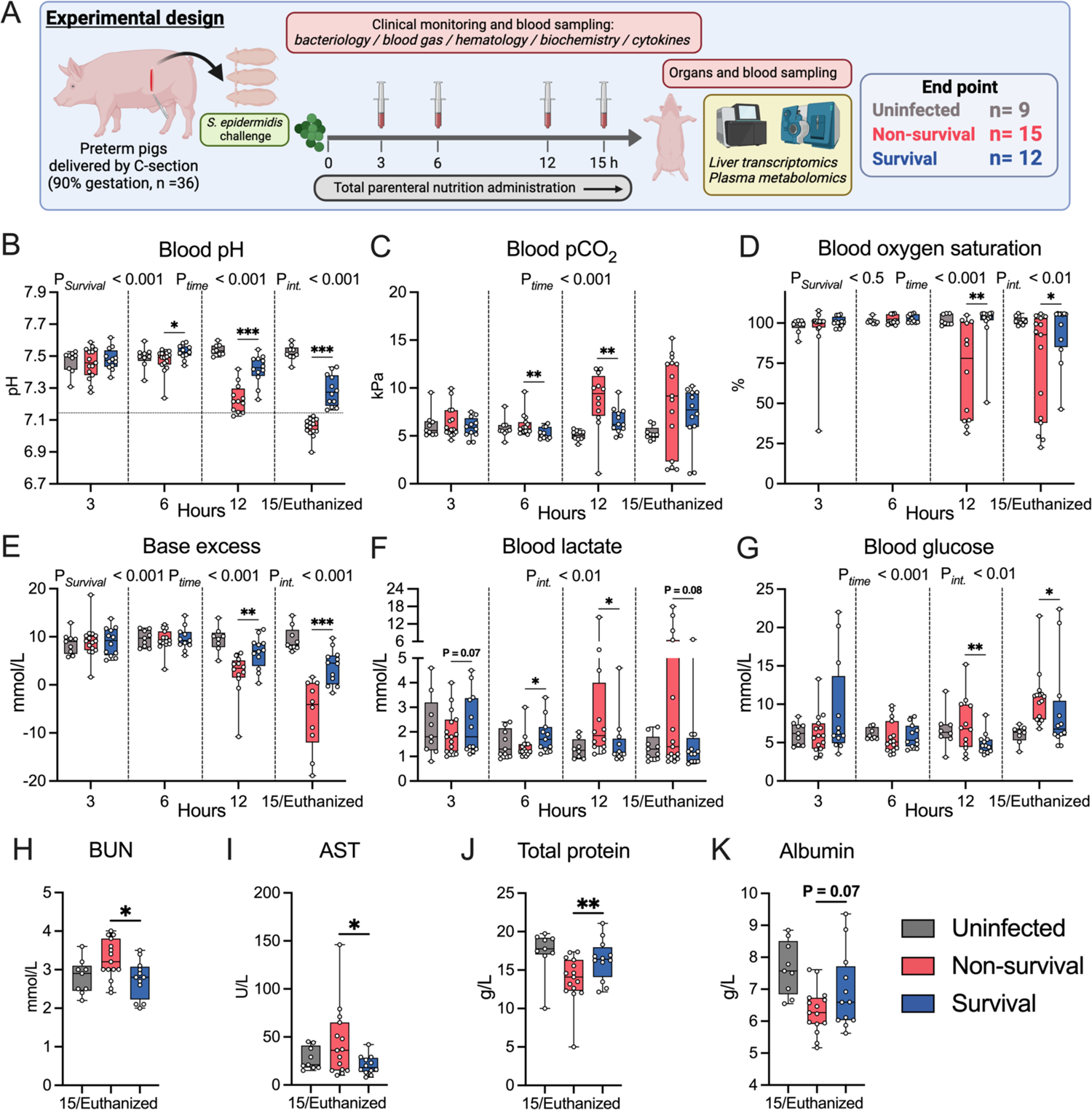
Clinical response disparities between infected survivors and non-survivors. Schematic of experimental design. Thirty-six cesarean-delivered preterm piglets were randomly assigned to receive either live *S. epidermidis* (10^9^ CFU/kg, n=27) or saline (n = 9), were nourished with total parenteral nutrition and monitored for 15 hours. This setup allowed for the division of piglets into three groups: survival (n = 12), non-survival (n = 15), and uninfected (n = 9). **(B-G)** Blood gas parameters (blood pH, blood pCO_2_, blood oxygen saturation, blood base excess, blood lactate, blood glucose) collected at 3, 6, 12, and 15 hours post-infection or euthanasia. The dotted horizontal line in the blood pH plot specifies the principal criterion for humane euthanasia. **(H-K)** Plasma biochemistry parameters (blood urea nitrogen (BUN), aspartate transaminase (AST), total protein, and albumin) collected at 15 hours post-infection or euthanasia **(B-K)** Data at each time point were analyzed separately via a linear mixed-effects model, incorporating group, gender, and birth weight as fixed factors and litter as a random factor. *P < 0.05, **P < 0.01, and ***P < 0.001, compared between survivors and non-survivors at the same time point. **(B-G)** Another linear mixed-effects model was employed to probe further disparities spanning the entire experimental duration, incorporating group, time, their interaction, gender, and birth weight as fixed factors, with litter and pig ID as random factors. P*_Survival_*, P*_time_*, and P*_int._* denote probability values for group effect (Survival vs. Non-survival) over time, time effects, and the interaction effects between time and group in the linear mixed effects interaction model, respectively. Uninfected animals served as a reference and were not included in the statistics. Statistical significance was defined as P-value < 0.05. All data were presented as box and whisker plots showing the range from minimum to maximum values. Panel A was created using Biorender.com.

Clinical parameters from blood gas analyses revealed distinct trajectories between survivors and non-survivors. Survivors showed relatively stable status across the first 12 h, followed by a mild drop of pH, oxygen saturation, and base excess, whereas non-survivors displayed a tendency of respiratory acidosis from 12 h (Fig. 1B-E). Interestingly, blood lactate levels showed a mild increase in survivors at the early stages of infection (3-6 h, 1-3 mmol/L, Fig. 1F), suggesting early enhancement of aerobic glycolysis to provide energy for inflammatory responses to eliminate circulating bacteria. However, this was reversed after 12-15 h as non-survivors started to develop metabolic acidosis, characterized by lactate levels > 5 mmol/L in several animals. Likewise, glucose homeostatic control was disrupted in non-survivors, as many non-survivors showed blood glucose levels > 10 mmol/L (Fig. 1G). Importantly, non-survivors also displayed signs of multiple organ injuries at the endpoint, including elevated levels of blood urea nitrogen and liver enzyme aspartate aminotransferase (indicating kidney and liver failure, respectively), as well as a significant drop of total blood protein and albumin levels (indicating capillary leakage) (Fig. 1H-K). These altered parameters are all classical signs of organ dysfunction in septic infants (18, 19). In summary, survivors showed relatively stable status during the infection, whereas infected non-survivors showed clear signs of both respiratory and metabolic acidosis during the early stages of infection, accompanied by vascular, kidney, and liver dysfunctions at the endpoint.

### Early enhanced resistance and later stronger tolerance responses contribute to infection survival

Next, we aimed to quantify the bacterial burden and cytokine levels in blood over the course of infection. In line with the early enhanced lactate production, infected survivors showed lower bacterial burdens (Fig. 2A) and reduced anti-inflammatory response (IL-10, Fig. 2G) at 3 h post-infectious challenge, compared to non-survivors. This further indicates an enhanced early resistance response, actively eliminating bacteria with an increased aerobic glycolysis rate, leading to lactate production despite an adequate oxygen supply (Fig 1D). Likewise, survivors showed higher levels of major leucocyte subsets at 3 and/or 6 h post-infectious challenge (Fig 2B-D). However, from 12h post-infectious challenge, survivors showed lower levels of lymphocytes (Fig 2B-D), with a decreased production of pro-inflammatory cytokine than non-survivors (IL-6, Fig. 2F), but a similar level of anti-inflammatory cytokines (IL-10, Fig. 2G), seemingly preventing a hyper-inflammatory response (Ratio of TNF-α/IL-10 and IL-6/IL-10, Fig. 2H-I).

**Fig. 2:**
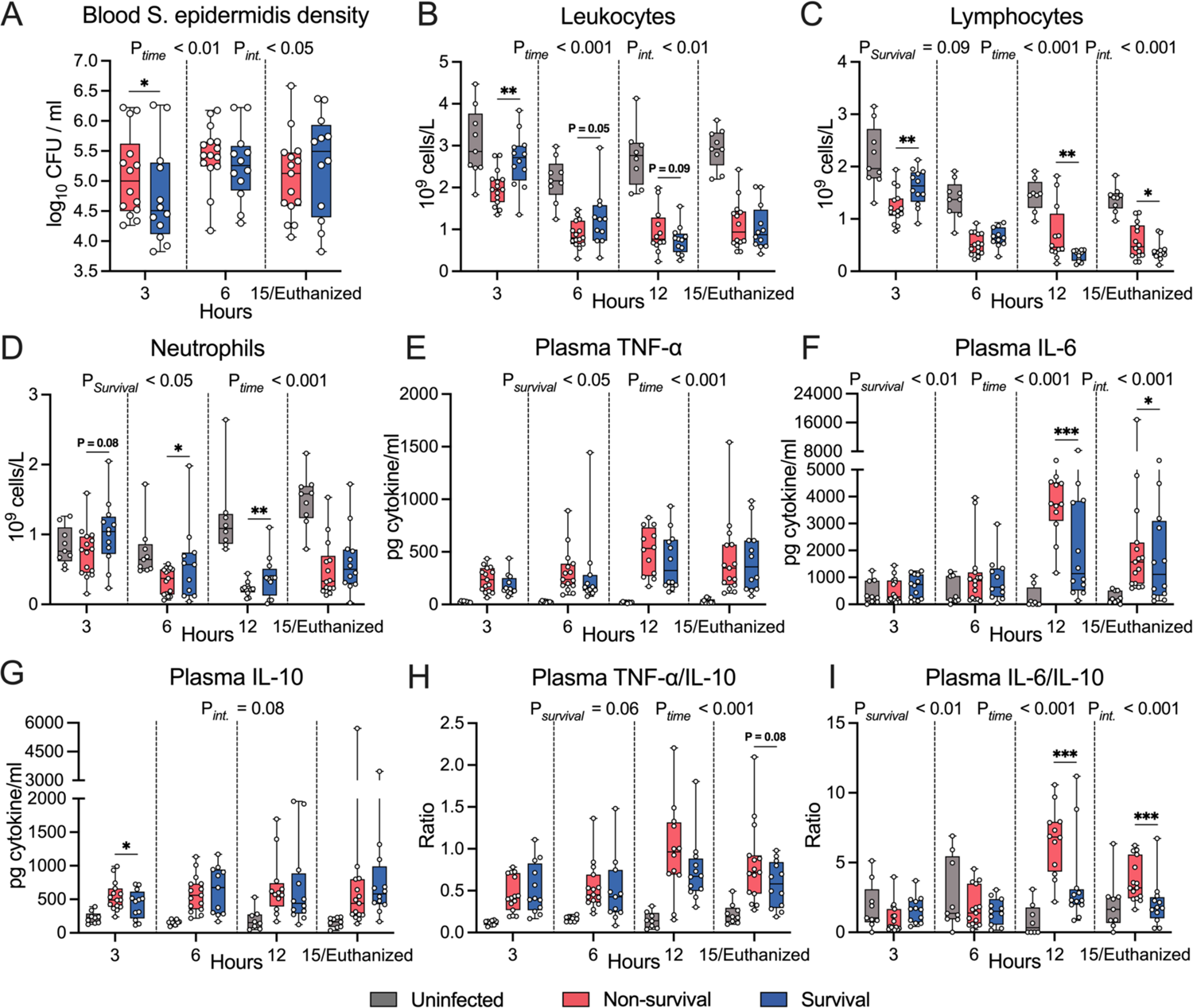
Bacterial burden, major leucocyte subsets, and cytokine levels in blood over the course of infection. The density of *S. epidermidis* in blood samples collected via jugular venous puncture at 3 and 6 hours or heart puncture at 15 hours, was then determined by counting colony-forming units (CFUs) after plating on tryptic soy agar containing 5% sheep’s blood and incubating for 24 hours at 37°C. **(B-E)** Numbers of major leukocyte subsets in blood samples collected 3-15 hours after *S. epidermidis* inoculation. **(E-G)** Cytokine levels were measured in heparinized plasma collected at 3, 6, 12, and 15 hours post-infection or euthanasia. **(H-I)** The ratio of TNF-α/IL-10 and IL-6/IL- 10, presented as 95% box plots **(A-I)** Data at each time point were analyzed separately via a linear mixed-effects model, incorporating group, gender, and birth weight as fixed factors and litter as a random factor. *P < 0.05, **P < 0.01, and ***P < 0.001, compared between survivors and non- survivors at the same time point. Another linear mixed-effects model was employed to probe further disparities spanning the entire experimental duration, incorporating group, time, their interaction, gender, and birth weight as fixed factors, with litter and pig ID as random factors. P*_Survival_*, P*_time_*, and P*_int._* denote probability values for group effect (Survival vs. Non-survival) over time, time effects, and the interaction effects between time and group in the linear mixed effects interaction model, respectively. Uninfected animals served as a reference and were not included in the statistics. Statistical significance was defined as P-value < 0.05. All data were presented as box and whisker plots showing the range from minimum to maximum values.

To investigate different strategies throughout the infection stages of sepsis survivors and non-survivors, we further explored the correlations between bacterial burdens and blood pH as well as inflammatory status at 3, 6, and 15 hours post-infection via a reaction norm analysis (7, 8, 20) (Fig. 3 & Fig. S1). First, we generated reaction norms using blood pH as our readout for general health, plotted against the pathogen burdens at each time point. A shallow slope indicates stable blood pH (health maintenance) despite a rising pathogen load, revealing a tolerance response. We found that both survivors and non-survivors exhibited similar slopes at 3 hours (Fig. 3A, left), together with previously described differences in bacterial burdens and major leucocyte subsets (Fig. 2A-D), suggesting enhanced resistance but not altered tolerance phenotype in infected survivors than non-survivors. This is supported by reaction norms of pro-inflammatory cytokines IL-6 and bacterial load with steeper slopes in survivors at 3 hours (Fig. 3B, left), implying augmented inflammation during the process of bacterial clearance. Importantly, at 6 h, infected survivors maintained a shallower slope than non-survivors in the blood pH-bacterial burden plot without distinct differences in cytokine-bacteria burden (Fig. 3A-B, middle & Fig. S1B, middle), indicating a stronger tolerance response in survivors. At the end of the study of 15 h, a clear distinction in blood pH-bacterial burden plot between the two groups was observed (Fig. 3A, right), suggesting survivors exhibited weaker tolerance than at 3 and 6 hours, while non-survivors showed dysregulated immune responses. Moreover, the less steep slopes in the cytokines-bacterial burden plots (for all three cytokines) further indicated that surviving animals adapted to the infection state to maintain vital organ functions rather than using their energy for inflammatory efforts (Fig. 3B, right, Fig. S1A-B, right).

**Fig. 3:**
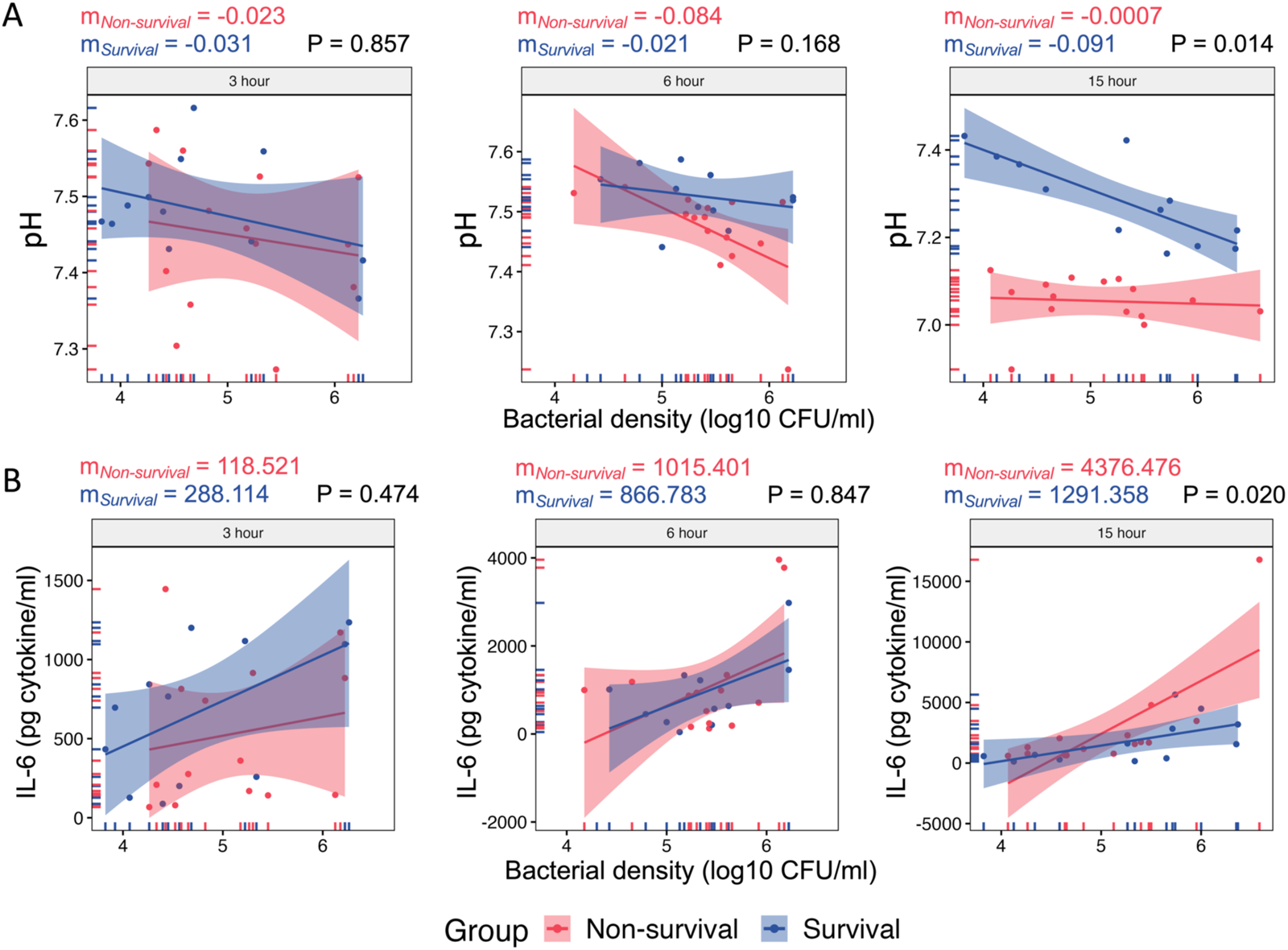
Early enhanced resistance and later stronger tolerance responses contribute to infection survival. **(A-B)** Reaction norm analysis was performed using blood pH (as a readout for health) and IL-6 (as a readout for inflammatory status) at 3, 6, and 15 hours, and plotted against the pathogen burdens at the same time point by linear regression, respectively. Extra sum-of-squares F Test was used to compare slopes. Statistical significance was defined as P-value < 0.05.

Collectively, we observed a delicate trajectory of different defense strategies in infected survivors, from exhibiting enhanced resistance at the early stage of infection to a stronger tolerance response and host-pathogen cooperation at the later stage of infection.

### Suppressed hepatic aerobic glycolysis but elevated mitochondrial activity in infected survivors

Clinical parameters, markers of organ dysfunctions, bacterial burdens, inflammatory profiles, and reaction norm analysis all suggest that enhanced tolerance but not resistance at the later phase of infection is a key driver of sepsis survival. Next, we aimed to investigate how alterations in the host metabolism regulate the induction of a tolerance phenotype to promote infection survival. As the first step, we performed bulk RNA sequencing (RNA-seq) of the liver tissue collected from animals belonging to the same litters (6 survivors, 10 non-survivors, and 6 uninfected piglets). Principal component analysis (PCA) revealed distinct gene expression profiles of uninfected animals, survivors, and non-survivors (Fig. 4A). To investigate the important pathways differently regulated across three groups, we employed a gene set enrichment analysis (GSEA) with the KEGG database. Compared to non-survivors, survivors exhibited downregulation of various carbohydrate metabolism pathways (such as glycolysis/gluconeogenesis, galactose metabolism, and starch and sucrose metabolism) as well as multiple immune and inflammatory pathways (including HIF-1α, Th1/Th2, Th17, IL-17 signaling, and various innate immune signaling pathways (Fig. 4B, left, Table S1A). In contrast, survivors showed an upregulation of pathways related to mitochondrial function, such as OXPHOS and nicotinate and nicotinamide metabolism. Subsequently, we extracted those differentially regulated pathways identified in survivors vs. uninfected and non-survivors vs. uninfected (Fig. 4B, middle and right, Table S1B-C). It was observed that both survivors and non-survivors exhibited similar enriched pathways when compared to uninfected individuals, with downregulation of OXPHOS and upregulation of multiple immune and inflammatory pathways. Furthermore, when validating the enriched pathways between survivors and non-survivors using GSEA with the GO:BP database, we observed a downregulation of several carbohydrate metabolism pathways (including pyruvate metabolic process, glycolytic process, and glucose metabolic process) in infected survivors relative to non-survivors (Fig. S2A, Table S1D). Conversely, an upregulation of ATP synthesis coupled electron transport, OXPHOS, mitochondrial respiratory chain complex assembly, and aerobic electron transport chain were upregulated in infected survivors.

**Fig. 4:**
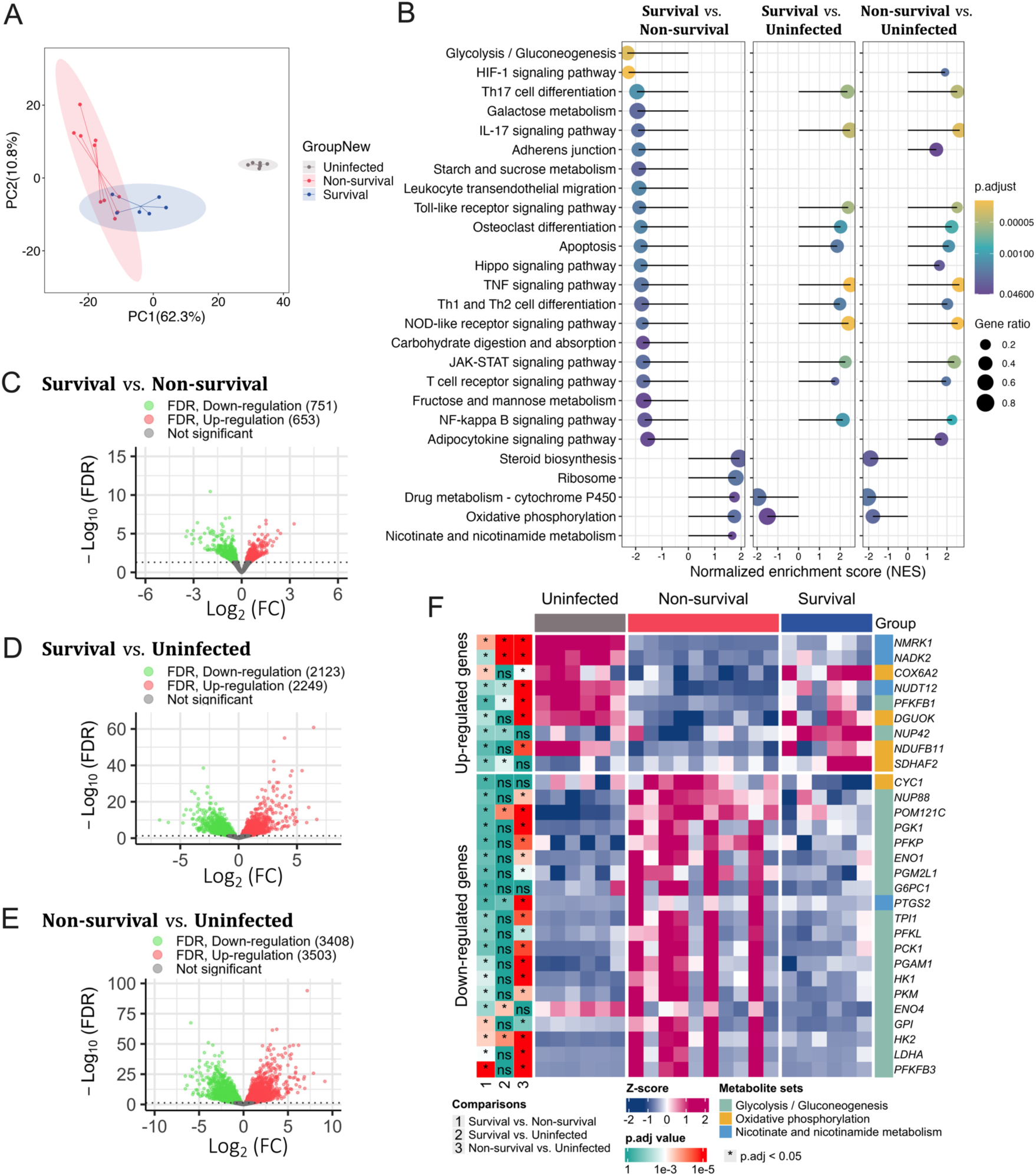
Suppressed hepatic aerobic glycolysis but elevated mitochondrial activity in infected survivors. **(A)** Principal component analysis (PCA) based on all annotated genes. Scores of the first two principal components from PCA were plotted. **(B)** A gene set enrichment analysis (GSEA) utilizing the KEGG database for *sus scrofa* (pig) was performed across three groups, and top-related enriched pathways with significantly expressed categories (adjusted P-value < 0.05 (FDR)) were presented. The size of the dots indicates the gene ratio, while the yellow color indicates a lower adjusted P-value. **(C-E)** Volcano plots between survivors vs. non-survivors, survivors vs. uninfected, and non-survivors vs. uninfected. A threshold of adjusted P-value < 0.05 (FDR) was set to illustrate the DEGs. **(F)** Heatmaps illustrating 29 DEGs involved in the top enriched pathways between survivors and non-survivors. The label of * represents adjusted P-value < 0.05 (FDR).

We then examined differentially expressed genes (DEGs) across three groups: survivors vs. non-survivors, survivors vs. uninfected, and non-survivors vs. uninfected. Our analysis revealed that survivors exhibited 1404 DEGs (653 upregulated and 751 downregulated genes) relative to non-surviving piglets. In contrast, when compared to uninfected piglets, survivors showed 4372 DEGs (2249 upregulated and 2123 downregulated), while the non-survivors demonstrated a significant difference with 6911 DEGs (3503 upregulated and 3408 downregulated) (Fig. 2C-E). Based on the enriched pathways from GSEA, we selected a set of 29 DEGs between surviving and non-surviving piglets involved in glycolysis/gluconeogenesis, OXPHOS, and nicotinate and nicotinamide metabolism (Fig. 4C). Four-fifths of genes involved in OXPHOS (*COX6A2*, *DGUOK*, *NDUFB11*, and *SDHAF2*) and three-fourths of genes involved nicotinate and nicotinamide metabolism (*NMRK1*, *NADK2*, and *NUDT12*) were up-regulated in survivors vs. non-survivors. On the contrary, survivors exhibited downregulation of the majority of genes associated with glycolysis/gluconeogenesis. Notably, key genes involved in aerobic glycolysis, including *ENO1*, *ENO4, PGAM1*, *GPI*, *HK1*, *LDHA*, *PGK1*, *PKM*, and *TPI1*, were all downregulated in survivors. Also, when comparing the two infected groups with uninfected animals (Fig. 4C), the profile of 29 genes related to the 3 key metabolic pathways in survivors tended to be similar to that in uninfected animals. Taken together, the hepatic transcriptomic data suggests an association of increased mitochondrial activity and reduced glycolysis with increased tolerance and likelihood for survival.

### Plasma metabolome confirms metabolic regulations associated with tolerance phenotype

To further explore the metabolic signatures associated with survival, we performed untargeted global metabolic profiling in plasma collected from animals belonging to the same litters (6 survivors, 4 non-survivors, and 3 uninfected piglets). A total of 754 metabolites were annotated in both positive and negative modes. The application of quantitative filtering (relative standard deviation (RSD) >30%) reduced the number of annotated metabolites from 754 to 661. PCA analysis displayed a clear clustering among the three groups based on 661metabolites (Fig. 5A). Of 661 metabolites measured, 70 metabolites mapped to various metabolic pathways were observed to be significantly altered between survivors and non-survivors (P < 0.05, VIP score ≥ 1, Table S2A). Among the altered metabolites, 52 (74.3%) were up-regulated, and 18 (25.7%) were down-regulated in survivors compared to non-survivors. A pie chart (Fig. 5B) indicates that most altered metabolites belonged to the lipid (28.6%), amino acid (17.1%), and carbohydrate and energy (15.7%) pathways, followed by nucleotide (14.3%), xenobiotics (8.6%), and peptide (5.7%) metabolic pathways.

**Fig. 5:**
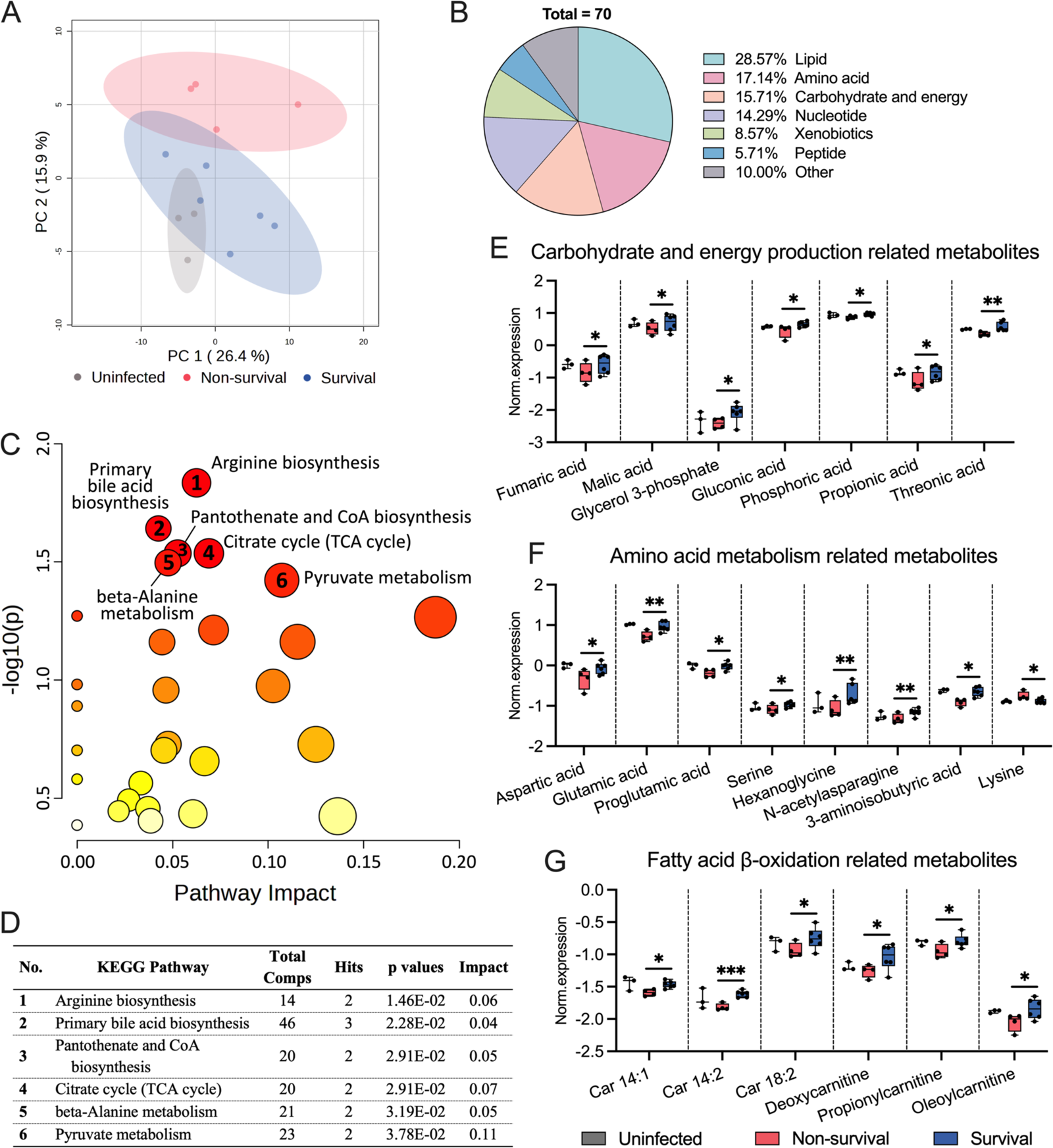
Plasma metabolome confirms metabolic regulations associated with tolerance phenotype. **(A)** Principal component analysis (PCA) based on 661 annotated metabolites from both positive and negative modes, with the scores of the first two principal components plotted to illustrate the group separation. **(B)** Pie chart depicting the classification of alerted metabolites between survivors and non-survivors. **(C)** Seventy metabolites with significant differences in abundance between survivors and non-survivors were subjected to the pathway enrichment analysis (y-axis for enrichment P-values, significance of a pathway) and the pathway topology analysis (x-axis for impact values, reflecting centrality and enrichment of a pathway) in the *MetaboAnalystR* package. The color of a circle indicates the level of enrichment significance, with red for high and yellow for low, and the size of a circle is proportional to the pathway’s impact value. **(D)** The table presents the top six pathways characterized by P-values with impacts, highlighting their significance between survivors vs. non-survivors. **(E-G)** Molecules with differential abundance in three main categories were presented as box and whisker plots showing the range from minimum to maximum values. A litter-adjusted linear model using the *limma* method was leveraged to identify MDAs with the P-value cut-off of 0.05 coupled with the variable of importance (VIP) score of the first component cut-off of 1. *P < 0.05 and **P < 0.01, compared between survivors and non-survivors at the same time point. Uninfected animals served as a reference and were not included in the statistics.

Feature-based pathway analysis was conducted using the *mummichog* algorithm, and 23 pathways were significantly enriched (Table S2B). These pathways were primarily associated with amino acid metabolism (D-amino acid metabolism; alanine, aspartate and glutamate metabolism; lysine metabolism; histidine metabolism; and phenylalanine, tyrosine and tryptophan biosynthesis), and carbohydrate metabolism (pentose phosphate pathway; pentose and glucuronate interconversions; and galactose metabolism). Lipid metabolism, including linoleic acid metabolism, biosynthesis of unsaturated fatty acids, and arachidonic acid metabolism, were also enriched. To validate and refine the findings from our exploratory feature-based pathway analysis, we executed a subsequent pathway analysis focusing on seventy molecules with differential abundance (MDA) (Table S2C). The top six pathways include arginine biosynthesis, primary bile acid biosynthesis, pantothenate and CoA biosynthesis, citrate cycle (TCA cycle), beta-alanine metabolism, and pyruvate metabolism (Fig 5C-D). Importantly, a majority of the identified pathways are interconnected and ultimately lead to the TCA cycle as a downstream metabolic pathway. Arginine and alanine are glucogenic amino acids and play a pivotal role in metabolism by entering the TCA cycle and transforming into intermediates that facilitate gluconeogenesis (21). Pantothenate is a crucial precursor for the biosynthesis of coenzyme A (CoA), which is essential for the conversion of pyruvate to acetyl-CoA, and also for the transformation of α-ketoglutarate to succinyl-CoA within the TCA cycle (22).

To explore specific metabolites involved in amino acid metabolism, carbohydrate metabolism and energy production, and fatty acid β-oxidation, box plots were generated to visualize MDAs across three groups (Fig. 5E-G). All metabolites related to carbohydrate metabolism and energy production were upregulated in infected survivors (Fig. 5E). Specifically, fumaric and malic acids are intermediates in the TCA cycle and play a crucial role in energy production (23). Glycerol 3-phosphate can either enter glycolysis and the TCA cycle or contribute to lipid biosynthesis (24). The majority of metabolites associated with amino acid metabolism exhibited upregulation, with the exception of lysine (Fig. 5F). Of note, aspartic acid, glutamic acid, and serine are all glucogenic amino acids that provide fuel for the TCA cycle (21). Furthermore, acylcarnitines, including Car 14:1, Car 14:2, Car 18:2, deoxycarnitine, propionylcarnitine, and oleoylcarnitine were all found to be upregulated in survivors (Fig. 5G). Acylcarnitines are involved in the transportation of fatty acids into the mitochondria for β-oxidation, which ultimately leads to the production of acetyl-CoA and provides energy for cell activities (25). Overall, our data suggests that the TCA cycle and a series of interconnected pathways are enhanced in survivors. This further strengthens hepatic transcriptomic findings of a late tolerance phenotype associated with rewired metabolic regulations to reduce glycolysis and prioritize mitochondrial activities as key mechanisms to survive the neonatal infection.

## DISCUSSION

Preterm infants, with lower energy reserves than term infants and adults, display unique defense strategies during infectious challenges, as energy is primarily devoted to essential growth and development (4). This allocation may result in a skewed defense tolerance phenotype, where tolerance acts as a protective mechanism in the early stages of infection by moderating excessive immune responses and subsequent tissue damage. However, resistance and tolerance are both crucial strategies in combating infections, each playing distinct roles in the host’s defense mechanisms against pathogens. By inhibiting pathogen growth and reproduction, resistance can decrease the transmission of the pathogen and, consequently, the prevalence of disease in the host (20). Nevertheless, if the resistance response is not properly controlled, it may lead to excessive inflammation, causing severe tissue damage and potentially sepsis. In this study, we suggest that triggering a resistance response through immune-metabolic regulators during early neonatal infection could prevent excessive bacterial growth in infected survivors. However, survivors are likely to exhibit an enhanced tolerance response, characterized by regulated inflammation and minimized tissue damage in the later stages of infection.

We have established a preterm piglet model of neonatal coagulase-negative staphylococci (CONS) sepsis induced by *S. epidermidis* infection with a consistent mortality rate of 40-50% during 15 h post-infectious challenge (14, 17). Septic non-survivors suffer from both respiratory and metabolic acidosis and vascular, kidney, and liver dysfunctions. Importantly, in this model, glucose supply and the degree of systemic glycolysis are the main drivers of disease outcomes (17). In the current study, we thereby asked why animals with similar genetic backgrounds and identical rearing regimes can display two distinct outcomes upon infection: survival or non-survival. We postulated that these two groups of animals may possess distinct host metabolism and defense strategies. We discovered that infected survivors showed stronger antimicrobial, glycolytic, and immune responses during the early phase of infection to control the prevalence of invading bacteria. This early enhancement of resistance was followed by a stronger tolerance phenotype during later infection, as demonstrated by similar bacterial burdens but better clinical status, glucose homeostasis, and immune and inflammatory control from 6-15 h post-infection in survivors compared to non-survivors.

Recognizing varying defense mechanisms throughout infection stages, from early enhanced resistance to later stronger tolerance in survivors, may have important implications for clinical practice and the development of novel therapeutic strategies. The current study shows that the increased resistance response during the first few hours post-challenge resulted in better clinical outcomes during later stages of infection, as demonstrated by reduced blood acidity and fewer respiratory difficulties in survivors than non-survivors. However, if an anti-inflammatory therapy is initiated in this critical early phase of infection, resistance response may be inhibited, potentially leading to bacterial overgrowth later and worsening clinical outcomes in the later phase. This has been observed in many previous studies in rodents (26–28) as well as in our own laboratory when we found that the anti-inflammatory inter alpha inhibitor protein (IAIP) at the early phase of infection exacerbated sepsis outcomes (14). Similar mechanisms may also explain why many randomized controlled trials testing immune suppressive sepsis therapies did not succeed (29–34).

Early resistance response observed in infected survivors in this study is associated with increased blood lactate (a marker of increased glycolysis), increased major leucocyte subsets levels, and diminished anti-inflammatory response (IL-10 levels). Immune effector cells, such as macrophages, exert resistance response via a metabolic switch from OXPHOS to aerobic glycolysis upon TLR stimulation and require a steady glycolytic flux to maintain the inflammatory phenotype (12). The most immediate consequences of aerobic glycolysis are increased intracellular and extracellular lactate concentrations (35). However, prolonged resistance phenotype causes significant risks of collateral tissue damage via unspecific immune responses. Therefore, maintenance of tolerance at the later phase of infection is crucial for host survival. In this study, from 6h to the end of the study, survivors clearly exhibited better clinical outcomes, less inflammatory response, and organ injuries. Evidence of enhanced tolerance in survivors at the euthanasia time point was strongly supported by liver transcriptomics and plasma metabolomics. In our hepatic transcriptomic analysis, we found a reduced expression of aerobic glycolysis-associated genes and several immune and inflammatory pathways in sepsis survivors, whereas OXPHOS and nicotinate and nicotinamide metabolism were upregulated in survivors. These regulations likely inhibit vast ATP production, which is necessary for strong immune responses, and instead support the maintenance of organ functions. Similar observations were shown previously in septic patients, demonstrating transcriptomic profiles with highly up-regulated glycolysis and HIF-1α signaling (36). When conducting plasma metabolomics, we found metabolic pathways related to the TCA cycle and mitochondrial activities were significantly enhanced in sepsis survivors. In a study involving over 1000 adult sepsis patients, metabolomics and proteomics analyses of plasma samples identified significant differences in fatty acid transport and β-oxidation, gluconeogenesis, and the TCA cycle between survivors and non-survivors, highlighting potential metabolic targets for improving patient outcomes (37). As such, therapeutics, macronutrients, or supplements that support these energy-generating metabolic pathways may be a viable option to reduce morbidity from neonatal sepsis.

Despite significant observations, there are certain limitations that need to be acknowledged. First, it was not possible to characterize host metabolism at several time points during the infection course to obtain a more comprehensive view of metabolic pathways connected to early resistance response in survivors. Second, acknowledging the advantage of using pigs with a more similar immune system to that in humans (compared to rodents), crossbred preterm piglets do not have an identical genetic background. Therefore, genetic differences may contribute to sepsis survival. Lastly, our preterm sepsis model mimics neonatal CONS infection in preterm infants, and it cannot be excluded that neonatal infection with other bacterial species may exhibit different patterns of defense strategies between survivors and non-survivors.

In conclusion, our findings demonstrate potential immune-metabolic mechanisms regulating distinct trajectories of defense strategies to survive neonatal infection. This study emphasizes the importance of the relationship between early resistance response and later tolerance associated with prioritized host metabolism on mitochondrial activities. The study also highlights the potential of using infection therapeutics targeting host metabolism to improve tolerance. However, the time for therapy initiation needs to be carefully considered to avoid interference with early resistance response.

## Methods

### *S. epidermidis* culture preparation

Originating from frozen stock, 500 μL of the *S. epidermidis* bacteria (WT-1457, isolated from a septic infant) were cultured in 30 mL of tryptic soy broth for 17 hours at 37°C and 200 rpm. The OD of the culture was measured using a spectrophotometer, and the bacterial concentration was calculated based on a previously established conversion factor correlating OD to CFU. The culture was then centrifuged for 20 minutes at 3000g, and the resulting bacterial pellet was resuspended in sterile physiological saline at a concentration of 3 × 10^8^ CFU/ml. To confirm the actual concentration used in each experiment, the bacterial culture was plated onto tryptic soy agar and incubated overnight at 37°C.

### Animal experimental procedures and intervention

We collected a cohort of thirty-six crossbred preterm piglets (Landrace x Large White x Duroc), born by cesarean section at day 106 (90% gestation, term at day 117) from five healthy pregnant sows. After delivery, the newborn piglets were relocated to the intensive care unit, where they were settled in infant incubators preset to a temperature of 37 °C and supplemented with an additional oxygen flow of 1-2 liters per minute. When needed, animals were resuscitated by Doxapram (0.1 ml/kg im.), Flumazenil (0.1 ml/kg im.), tactile stimulation, and positive airway pressure ventilation, applied as necessary until achieving respiratory stability. Furthermore, each piglet was equipped with a vascular catheter inserted into the dorsal aorta via the transected umbilical cord, facilitating both the administration of parenteral nutrition (PN), bacterial inoculation, and blood sampling.

Approximately two hours postpartum, all animals received PN, and were categorized based on birth weight and gender, and then randomized to either receive live *S. epidermidis* (10^9^ CFU/kg, n = 27) or a control saline (n = 9) via an interatrial infusion over 3 minutes. Thereafter, the animals were intensively cared for until the 15-h post-infectious challenge. During this period, animals reaching predefined humane endpoint (sepsis criteria, including arterial blood pH of ≤ 7.1 and clinical signs of deep lethargy, discoloration, and tachypnea) were euthanized for tissue collection and counted as an episode of mortality (non-survival). Blood samples were collected at 3, 6, 12, and 15 h post-bacterial challenge via the arterial catheter for blood gas analysis and hematology, as well as plasma storage for cytokine measurements and metabolomic analyses. For the specific purpose of bacterial enumeration within the blood, samples were drawn through puncture of the jugular vein at the 3, 6, and 15-h time points. Serum samples at euthanasia were used for biochemical analysis.

### Euthanasia and organ collection

At humane endpoint or 15 h of post-infection, all piglets were deeply anesthetized with a Zoletil mixture (0.1 ml/kg), which included Zoletil 50 (125 mg tiletamine, 125 mg zolazepam), xylazine (6.25 ml xylazine 20 mg/ml), ketamine (1.25 ml ketamine 100 mg/ml), and butorphanol (2.5 ml butorphanol 10 mg/ml), and were subsequently euthanized with an intracardiac injection of barbiturate. The liver and spleen were collected and preserved (frozen and fixed) during necropsy.

### Blood gas, hematology, inflammatory markers, and serum biochemistry

Blood samples at 3, 6, 12, and 15 h underwent routine blood gas analysis utilizing the GEM Premier 3000 (Instrumentation Laboratory, USA). Hematological assessments were conducted with the ADVIA 2120i Hematology System (Siemens, Germany). For the quantification of plasma cytokines, TNF-α, IL-6, and IL-10 were analyzed using porcine-specific DuoSet enzyme-linked immunosorbent assays (R&D Systems). Additionally, serum biochemistry evaluations were executed using the ADVIA 1800 Chemistry System (Siemens, Germany).

### Hepatic RNA-Seq transcriptome analysis

Liver RNA from all animals was extracted using the RNeasy mini kit (QIAGEN, US), and subsequently, whole-transcriptome shotgun sequencing was used for gene expression profiling. Library preparation and sequencing were performed by NOVOGENE services (Cambridge, UK). In brief, RNA-Seq libraries were prepared using 1000 ng of RNA and the VAHTS mRNA-Seq V3 Library Prep Kit for Illumina (Vazyme, Nanjing, PRC). The libraries were sequenced on the Illumina NovaSeq 6000 platform, generating 150 bp paired-end reads. Quality and adapter trimming of raw reads were performed using TrimGalore (Babraham Bioinformatics, Cambridge, UK). The resulting clean reads, approximately 26 million per sample, were aligned to the porcine genome (*Sscrofa11.1*) using Tophat2 (38). Gene annotation information for the porcine genome was sourced from *Ensembl* (release 99). The script htseq-count (39) was employed to generate a gene-count matrix.

For differentially expressed gene (DEG) analysis, we excluded weakly expressed genes and only used protein-coding genes for downstream analysis. The *DESeq2* package version 1.38.3 (40) was implemented to acquire differentially expressed genes (DEGs) in three comparisons: Survivors *vs.* non-survivors, survivors *vs.* uninfected, and non-survivors *vs*. uninfected. No more covariates were added to the model. Herein, the robust estimation of fold change (FC) using the *lfcShrink* function with *ashr* and false discovery rate (FDR) cut-off of 0.05 was used (41). The volcano plots of total genes were visualized via the *EnhancedVolcano* version 1.14.0 package (42). The gene set enrichment analysis (GSEA) was used to elucidate the significantly perturbed pathways through the *clusterProfilter* package (version 4.0.2) (43). The FC in expression (log scale) obtained from DEG analysis was used to rank the gene list before introducing them into pathway analyses. Two knowledgebases of the Kyoto Encyclopedia of Genes and Genomes (KEGG) for *Sus scrofa*, and the Biological Processes aspect of Gene Ontology (GO:BP) for *Sus scrofa* were utilized. The FDR cut-off of 0.05 was applied to identify significant pathways. A normalized enrichment score (NES) was obtained in pathway analysis, with either negative or positive values indicating the pathways being down- or up-regulated. The relative expression of pathway-enriched genes was visualized via heatmaps using the *ComplexHeatmap* package version 2.15.1 (44). The analysis of liver transcriptomics data was achieved through R 4.2.3 (45).

### Plasma metabolome analysis

Untargeted metabolomics analysis was performed using ultra-performance liquid chromatography-mass spectrometry (UPLC-MS) via service from Creative Proteomics (Shirley, NY, USA). In brief, the methodology commenced with thawing the samples, followed by transferring 100 µL of each into a 1.5 mL tube and adding 300 µL of pre-cooled methanol. After vortexing the samples for 30 s, they were stored at -20 °C for 1 h and then centrifuged at 9660 *rcf* and 4 °C for 15 min. The supernatant was subsequently transferred to new tubes, stored again at - 20 °C for 1 h, and centrifuged under the same conditions. Afterward, 120 µL of the supernatant and 3 µL of DL-o-Chlorophenylalanine (0.5 mg/mL) were prepared in a vial for the instrumental analysis. The analysis was conducted using an ACQUITY UPLC system (Waters, Milford, MA, USA) combined with a Q-Exactive-Plus Hybrid Quadrupole-Orbitrap mass-spectrometer (Thermo, Waltham, MA, USA). Column was ACQUITY UPLC HSS T3 column of 100 mm × 2.1 mm; 1.8 µm (Waters, Milford, MA, USA). Mobile phase A was water with 0.05% formic acid and mobile phase B was acetonitrile. A gradient condition was: 0-1 min, 5% B; 1-12.5 min, 5% to 95% B; 12.5-13.5 min, 95% B; 13.5-13.6 min, 95% to 5% B; 13.6-16 min, 5% B. A fixed flow rate of 0.3 mL/min was used, and column temperature was maintained at 40 °C. The sampler was set at 4 °C during the analysis. Mass spectrometry parameters in both positive (POS) and negative (NEG) ion modes included a heater temperature of 300 °C, sheath gas flow rate of 45 arb, auxiliary gas flow rate of 15 arb, sweep gas flow rate of 1 arb, spray voltage of 3.0 kV for POS and 3.2 kV for NEG, capillary temperature of 350 °C, and S-Lens RF levels of 30% for POS and 60% for NEG.

A total of 8516 features in both positive and negative modes were detected, and 754 molecules were annotated (assigned by the vendor and cross-examined by our team). Molecule reproducibility was tested by calculating the variation of QC sample technical replicates (RSD, n = 6), and molecules with RSD >30% were discarded. For differential metabolites analysis, only annotated molecules were used for downstream analyses. To determine molecules with differential abundance (MDA), data first underwent sum normalization and log10 transformation without scaling. Subsequently, a litter-adjusted linear model using the *limma* method was leveraged to identify MDAs with the P-value cut-off of 0.05 coupled with the variable of importance (VIP) score of the first component cut-off of 1. Both feature-based functional analysis and MDA-based pathway analysis were leveraged to unveil perturbed pathways. For functional analysis, the mixed ion mode of POS and NEG created by the feature-wise merging of retention time, mass-to-charge values, and P-values obtained from statistical analyses of POS and NEG separately were used as input data. MDAs-based pathway analysis, which integrated Fisher’s exact test and out-degree centrality, was utilized. The knowledgebase from *Sus scrofa* of the Kyoto Encyclopedia of Genes and Genomes (KEGG) was used. Pathways with a P-value cut-off of 0.05 or a gamma-adjusted P-value cut-off of 0.05 and more than one significant hit with respect to functional and pathway analysis were further undergone biological interpretation. All analyses were conducted using *MetaboAnalystR* package version 4.0.0 (46) in R version 4.2.3 (45).

### Statistical analysis

All statistics were executed via R version 4.3.2 (45). For discerning differences at specific intervals (3, 6, 12, and 15 h), continuous data underwent analysis via a linear mixed-effects model, incorporating group, gender, and birth weight as fixed factors and litter as a random factor, using *lme4* package version 1.1.35.1 (47). Another linear mixed-effects model was employed to probe further disparities spanning the entire experimental duration. This model integrated group, time, their interaction, gender, and birth weight as fixed factors, with litter and pig ID as random factors, using *lme4* package version 1.1.35.1 (47). Normal distribution, variance homogeneity of residuals, and fitted values were assessed, and if the data did not conform to a normal distribution, logarithmic transformation was applied. If transformation did not achieve normality, the non-parametric Mann-Whitney U test was used instead. Uninfected animals served as a reference and were not included in the statistics. To examine different strategies in the infection response of sepsis survivors and non-survivors, reaction norm analysis was performed using blood pH (as a readout for health) and cytokines (as a readout for inflammatory status) at 3, 6, and 15 hours and plotted against the pathogen burdens at the same time point by linear regression. Extra sum-of-squares F Test was used to compare slopes. Statistical significance was defined as P-value < 0.05. Values for all data points in graphs are reported in the Supporting Data Values file.

## Supporting information

sFig.1-2, Table S1A-E, Table S2A-C

## Study approval

All animal studies and experimental procedures were approved by the Danish Animal Experiments Inspectorate under license number 2020-15-0201-00520. These approvals are in accordance with the EU Directive 2010/63, which governs the legislation for the use of animals in research.

## Data availability

The RNA-seq data generated from this study is deposited in the NIH Gene Expression Omnibus. The accession number for the raw and processed data is XXX (already uploaded, waiting for revision). The original and normalized metabolomics data were attached to the supplementary data. This study does not report custom computer code. Any additional information required to reanalyze the data reported in this paper is available from the lead contact upon request.

## Author contributions

DNN, OB, ZY, and JR designed the experiments. ZW, JR, OB, and DNN performed the animal experiments. ZY and JR performed laboratory analyses. ZY performed all statistical analyses of clinical data, managed raw data, and generated figures and tables. ZY, NT, and HY were responsible for all omics data analysis under DNN and NPL’s supervision. ZY, NT, OB, BK, NPL, and DNN were mainly responsible for data interpretation. ZY, NT, and DNN wrote the first draft of the manuscript. All authors contributed to data interpretation, manuscript revision, and approval of the final manuscript version. The levels of contribution in study design, actual experiments, laboratory analyses, and writing were used to assign co-first authorship order.

## Acknowledgments

The study was funded by the Novo Nordisk Foundation (grant number NNF220C0078747). ZY was supported by a PhD grant from the China Scholarship Council (CSC). The authors thank Tik Muk, Thomas Thymann, Simone Margaard Offersen, Anders Brunse, Karoline Aasmul-Olsen, Malene Spiegelhauer, Kristina Larsen, Malene Skovsted Cilieborg, Britta Karlsson, and Jane C. Povlsen for the assistance in animal experiments and lab analysis. Fig. 1A was created with Biorender.com.

## Notes

### Competing Interest Statement

The authors have declared no competing interest.

### Summary of Updates

author name correction (Tinh Thu Nguyen)

