## Supplementary material for "Regulation of host metabolism and defense strategies to survive neonatal infection": sFig.1-2, Table S1A-E, Table S2A-C

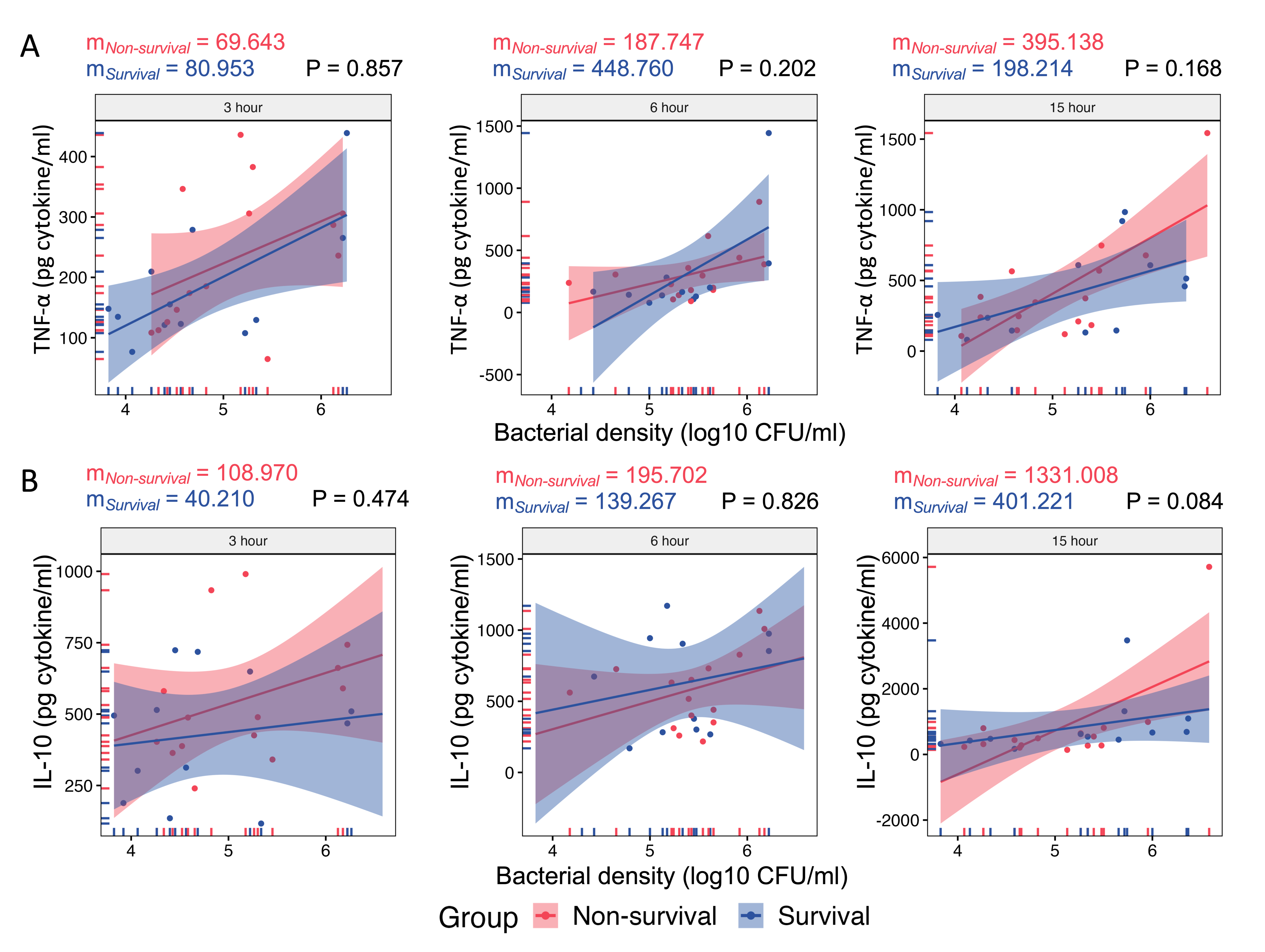

**sFig. 1: Reaction norms of cytokine-bacteria burden.**

**(A-B)** Reaction norm analysis was performed using blood pH (as a readout for health) and cytokines (TNF-α and IL-10) as a readout for inflammatory status at 3, 6, and 15 hours, and plotted against the pathogen burdens at the same time point by linear regression, respectively. Extra sum-of-squares F Test was used to compare slopes. Statistical significance was defined as P-value < 0.05.

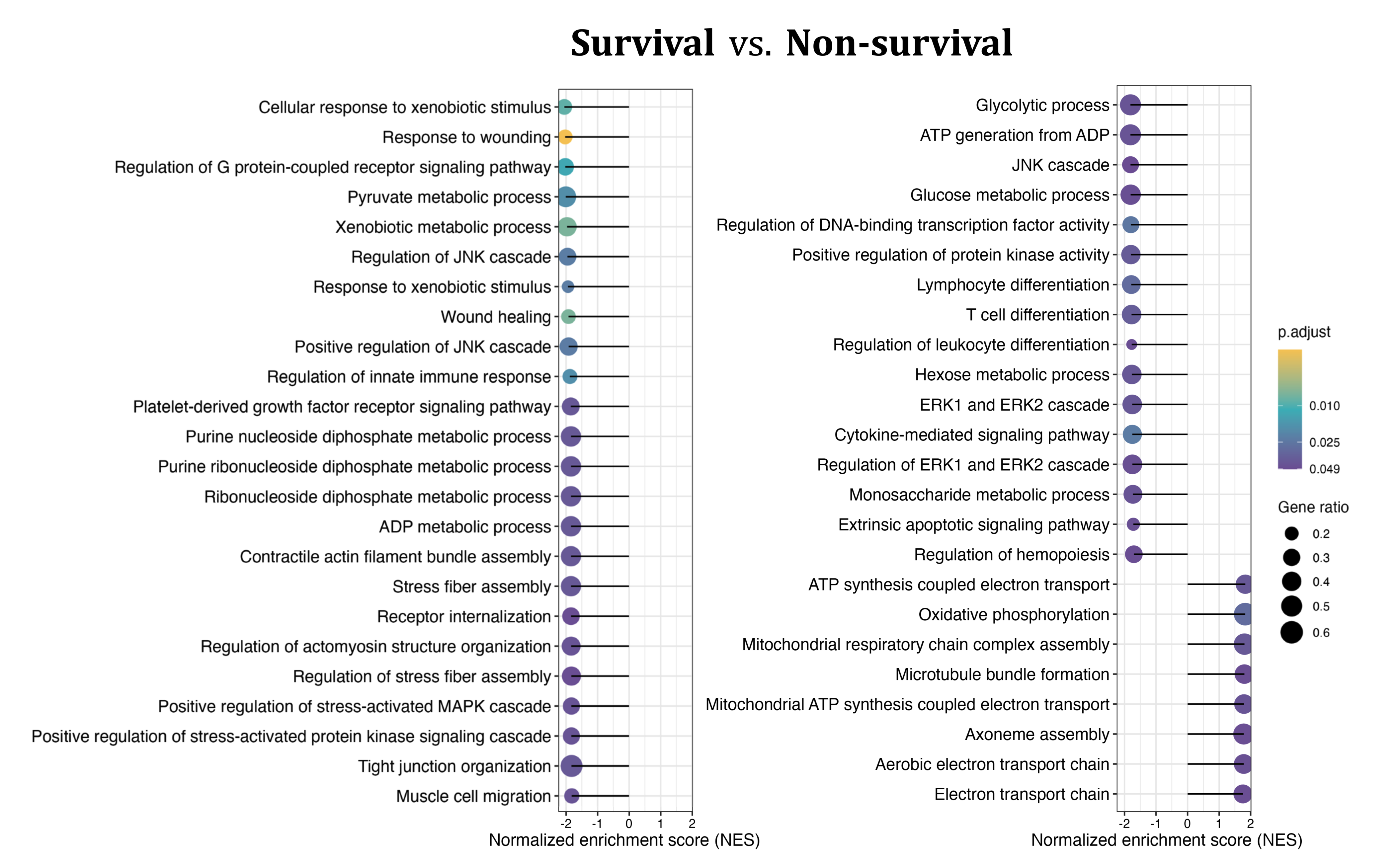

**sFig. 2:** **Suppressed hepatic aerobic glycolysis but elevated mitochondrial activity in infected survivors.**

A gene set enrichment analysis (GSEA) utilizing the GO:BP database for *sus scrofa* (pig) was performed to compare survivors and non-survivors, and top-related enriched pathways with significantly expressed categories (adjusted P-value < 0.05 (FDR)) were presented. The size of the dots indicates the gene ratio, while the yellow color indicates a lower adjusted P-value.

**Table S1A** KEGG-based gene set enrichment analysis result (Survivors vs. Non-survivors)

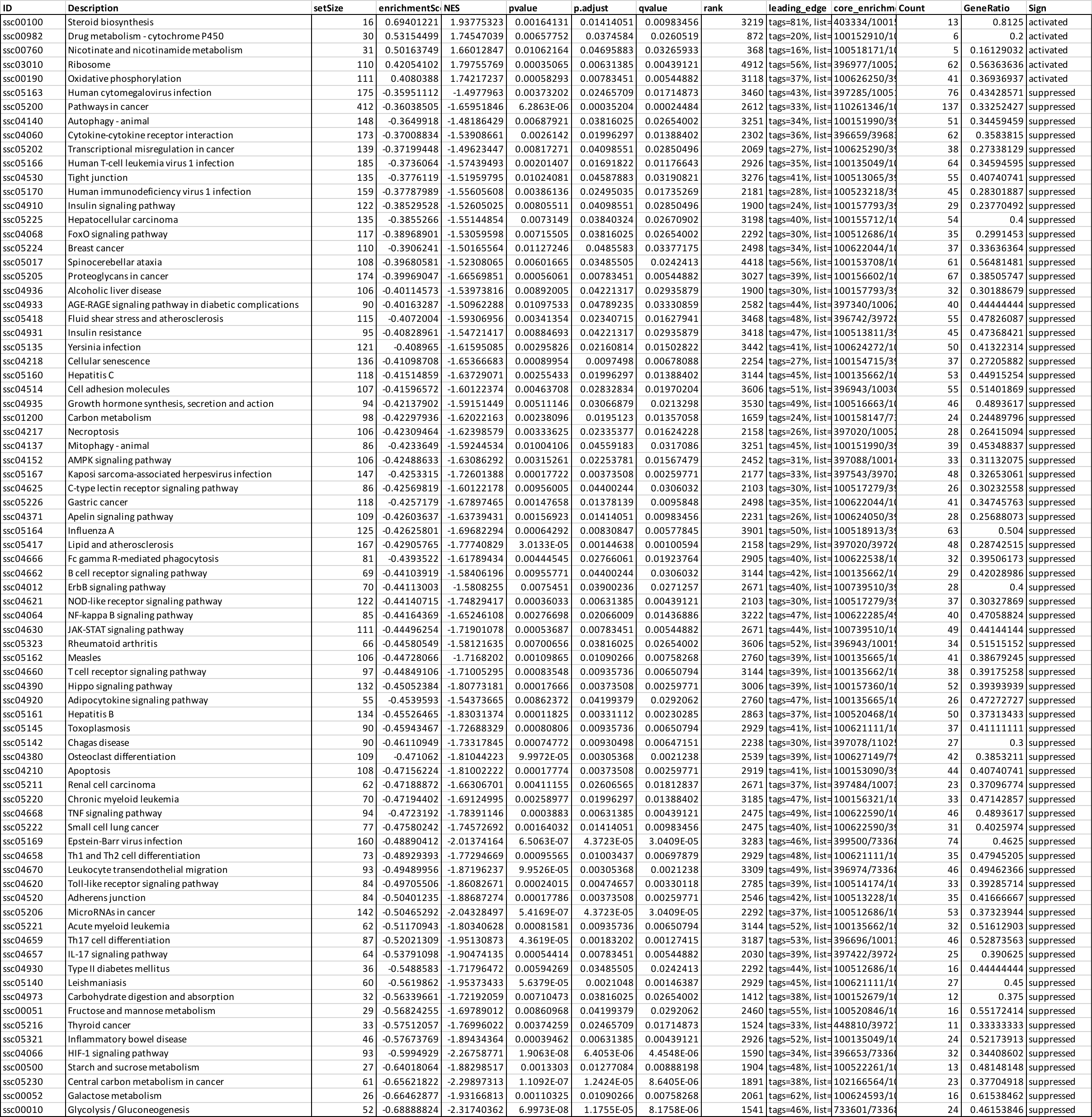

**Table S1B** KEGG-based gene set enrichment analysis result (Survivors vs. Uninfected)

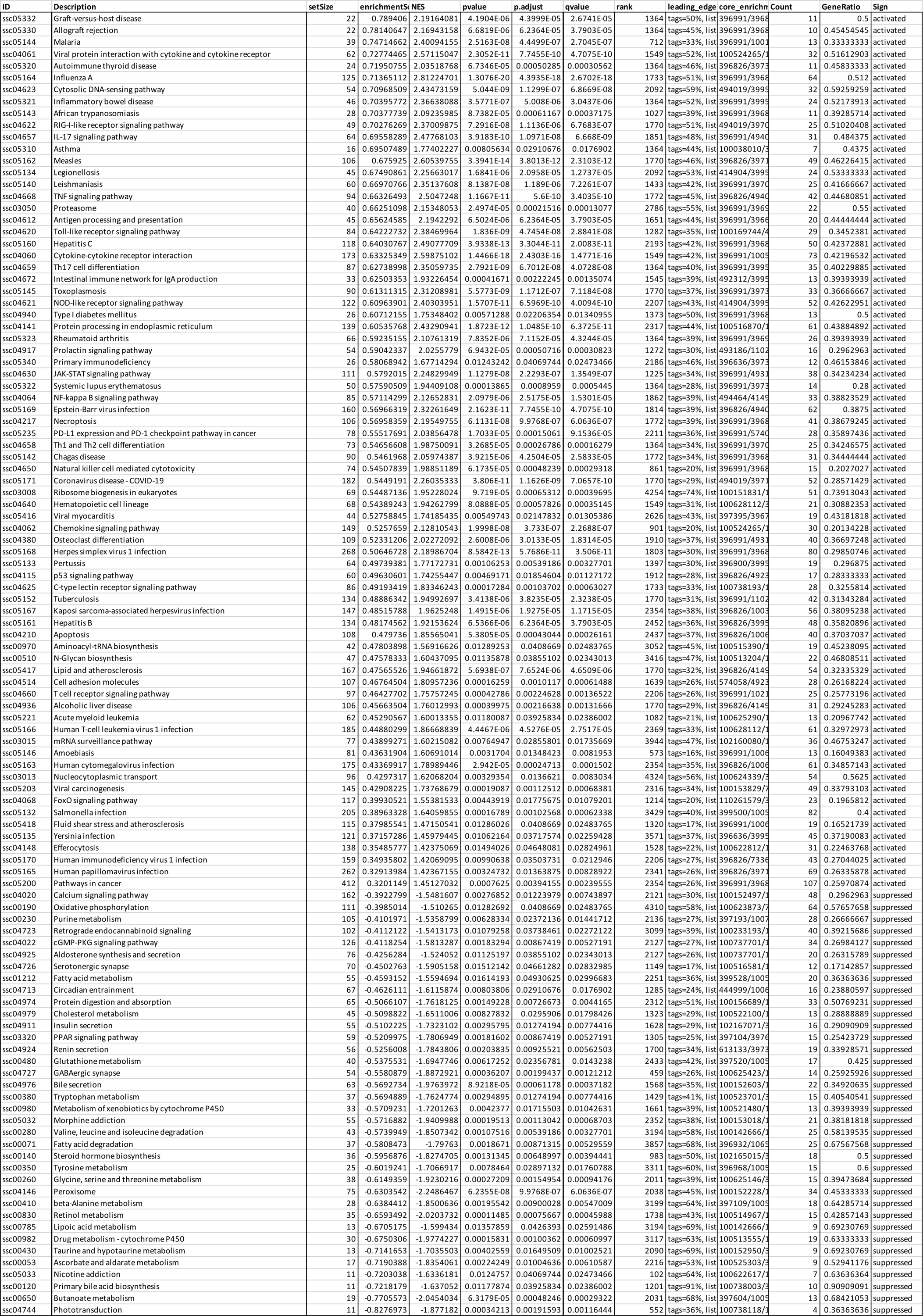

**Table S1C** KEGG-based gene set enrichment analysis result (Non-survivors vs. Uninfected)

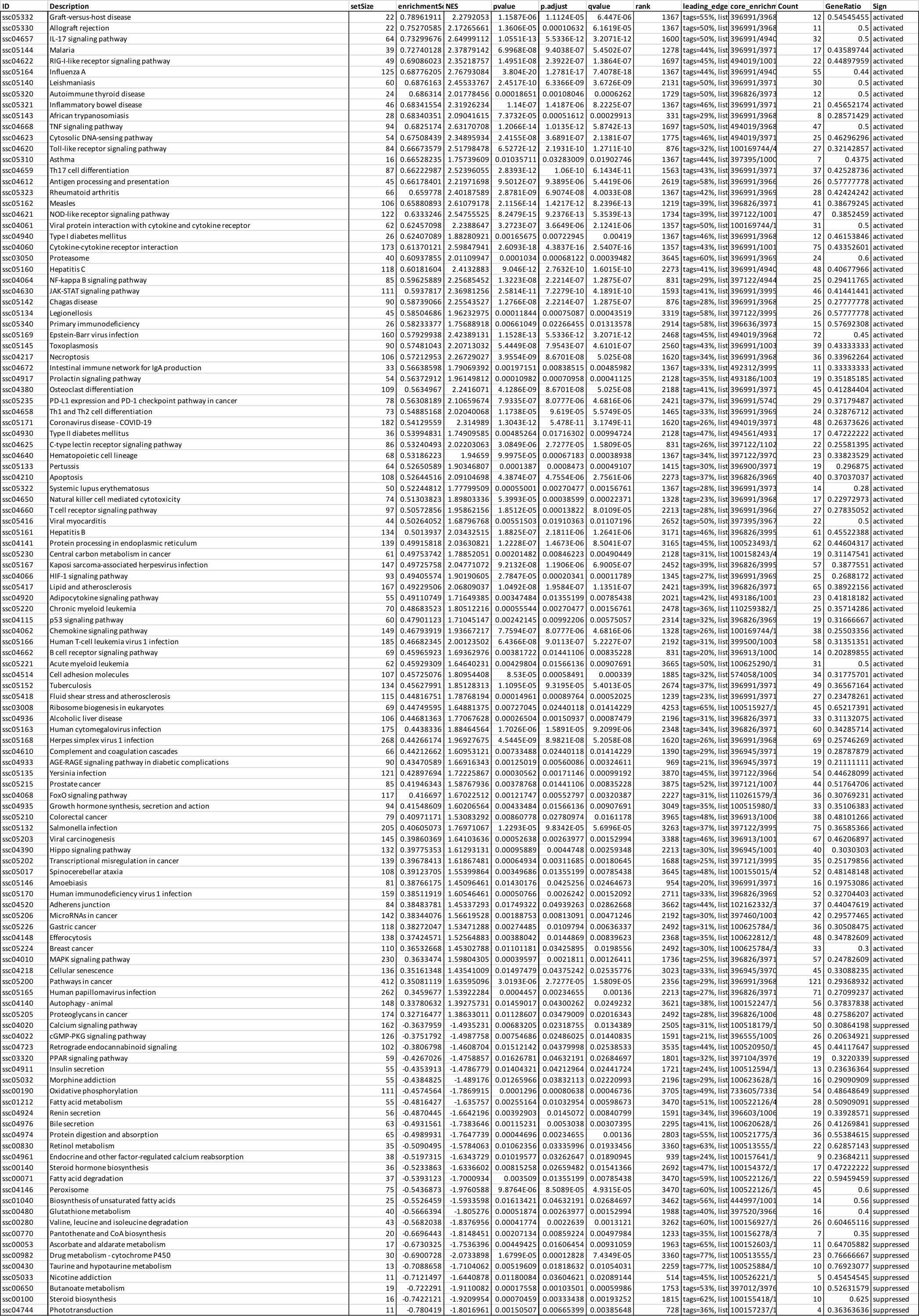

**Table S1D** GO:BP-based gene set enrichment analysis result (Survivors vs. Non-survivors)

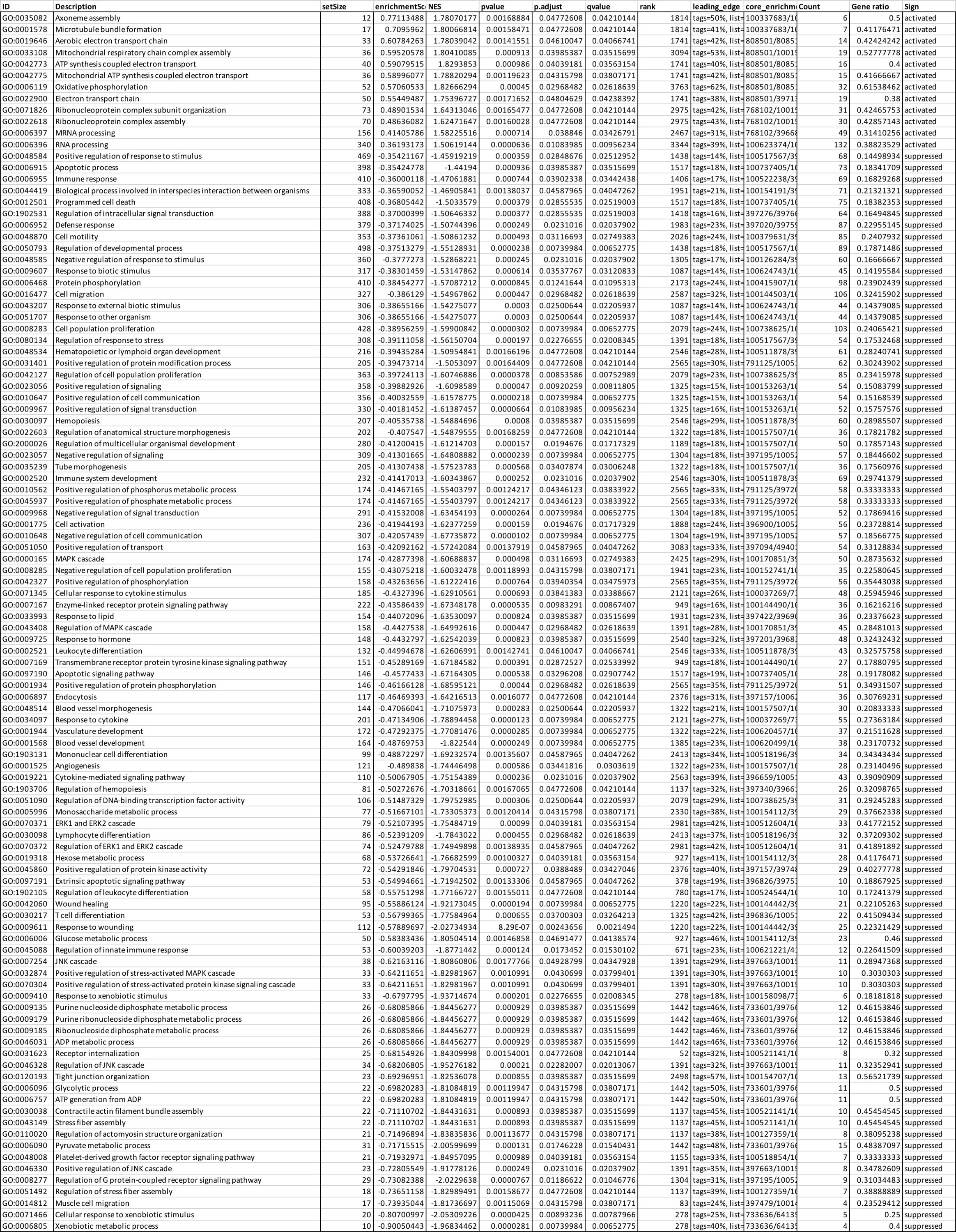

**Table S1E** Animal ID for NIH Gene Expression Omnibus identification

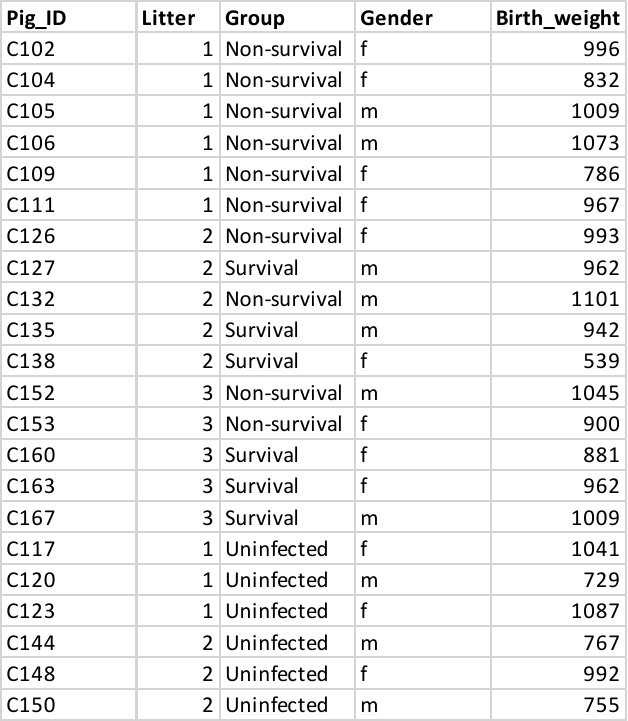

**Table S2A** Molecules with differential abundance

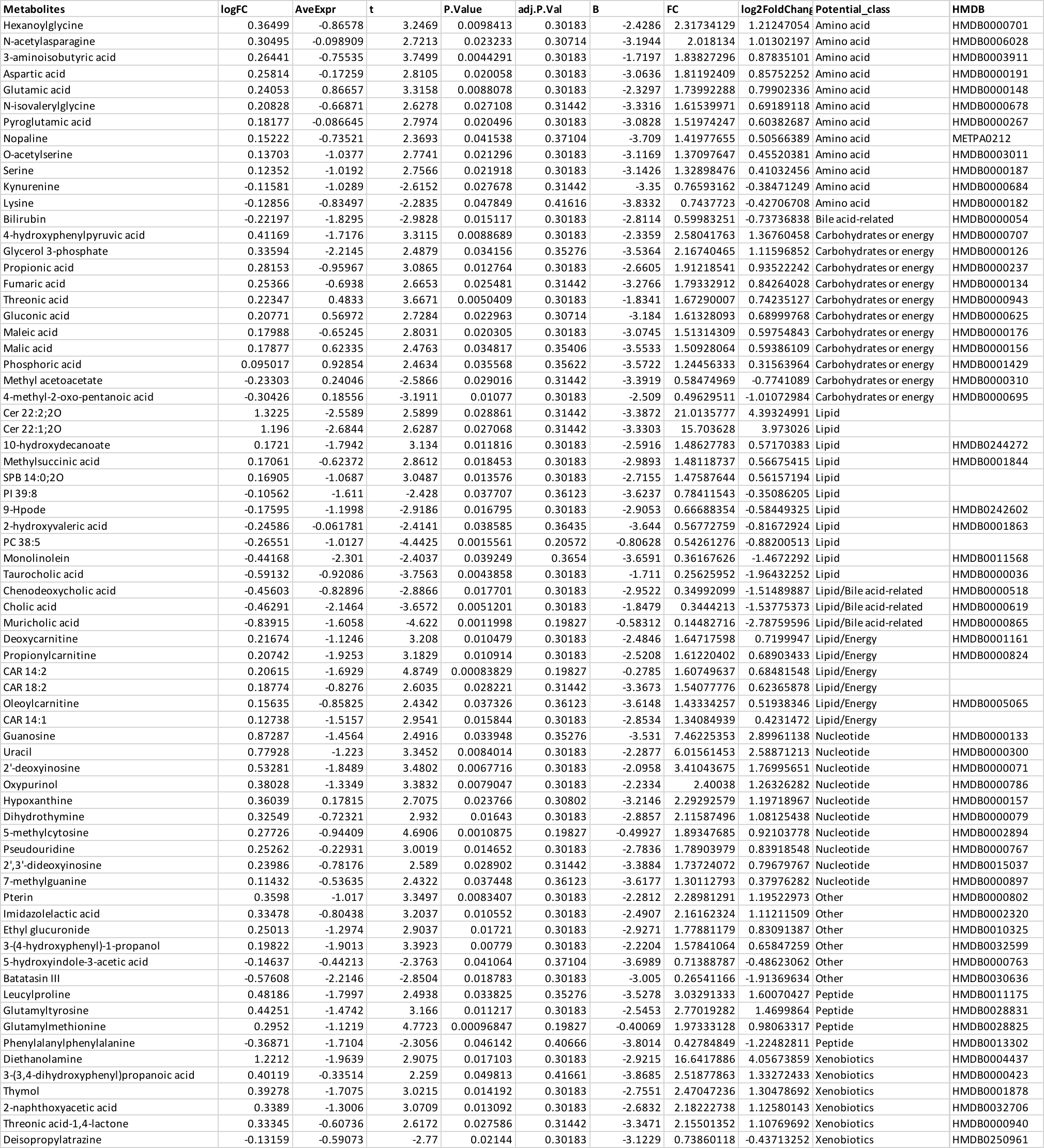

**Table S2B** Feature-based functional analysis result

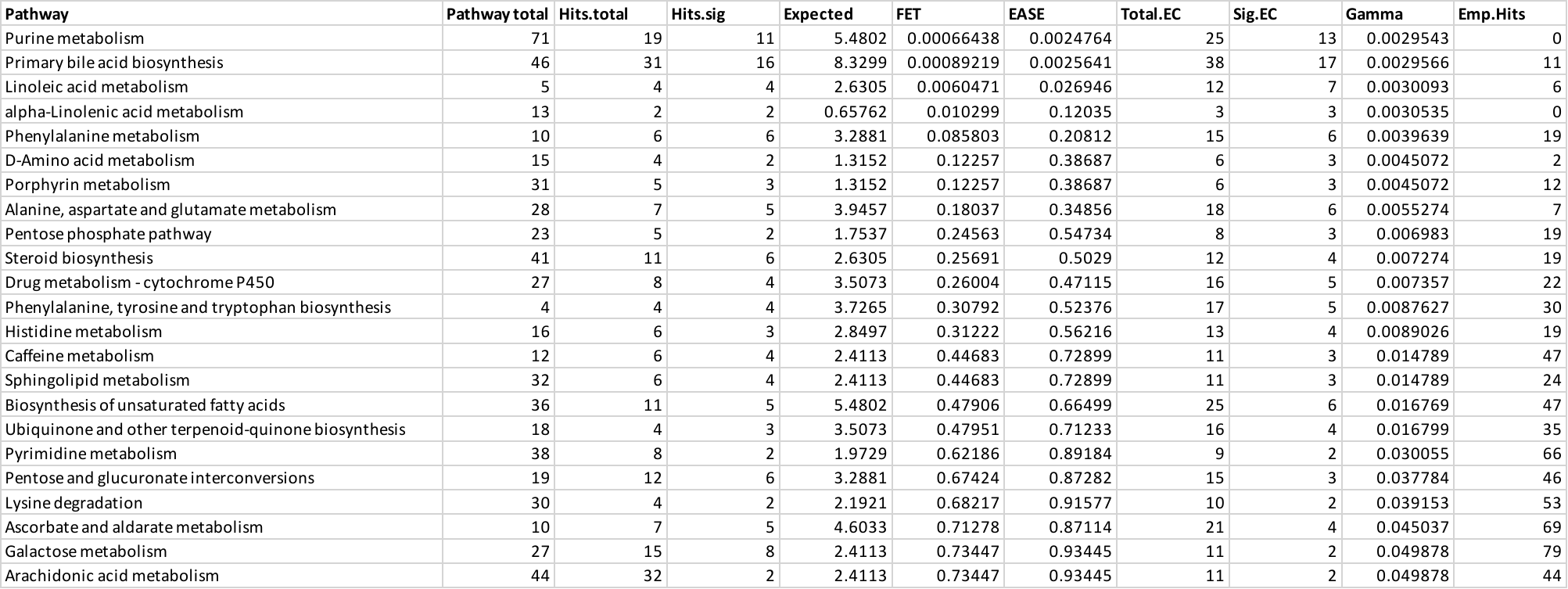

**Table S2C** MDAs-based pathway analysis result

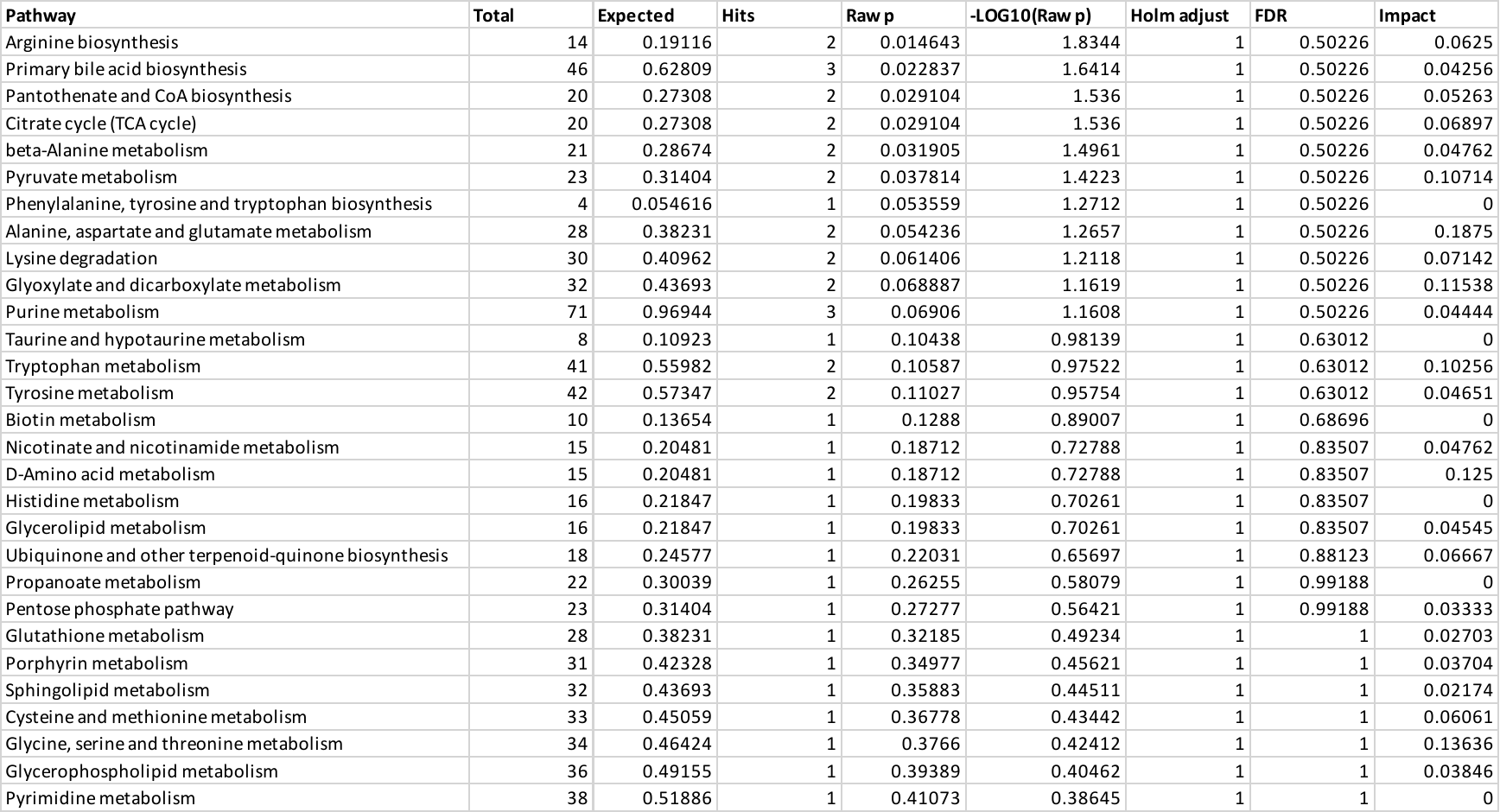
